# Super-resolution microscopy informs on the molecular architecture of alpha-synuclein inclusions in model systems and in the human brain

**DOI:** 10.1101/2021.04.25.441304

**Authors:** Patrick Weish, Diana F. Lázaro, Luís Palmares, Patrícia I. Santos, Christine Stadelmann, Günter U. Höglinger, Silvio O. Rizzoli, Tiago F. Outeiro

## Abstract

Lewy bodies (LBs) and Lewy neurites are pathological hallmarks of Parkinson’s disease and other progressive neurodegenerative disorders known as Lewy body diseases (LBD). These proteinaceous deposits are immunopositive for alpha-synuclein (aSyn) and several other proteins, as neurofilament components. The structural organization and composition of aSyn inclusions is still unclear and needs to be addressed in greater detail, as this may open novel avenues for our understanding of the disease-relevant pathological events.

In this study, we investigated the molecular architecture of aSyn inclusions, both in cell models and in human brain tissue, using state-of-art super resolution X10 Expansion microscopy (ExM). This approach physically expands specimens embedded into a swellable gel, preserving their biological information. Then, the specimen can be analyzed using standard epifluorescence microscopes, thereby obtaining nanoscale information.

The combination of different cell models, mouse and human brain tissue enabled us to distinguish different types aSyn assemblies (e.g. ring shape or tubular structures), and a conserved pattern of aSyn inclusions surrounded/encaged by intermediate filament proteins. Overall, X10 ExM enabled us to gain insight into the architecture and biology of aSyn inclusions and constitutes a powerful tool in the quest to understanding underlying disease mechanisms in synucleinopathies.

## Introduction

The accumulation of misfolded proteins in the brain is a hallmark shared by several neurodegenerative diseases, like Parkinson’s disease (PD), and dementia with Lewy bodies (DLB). The accumulation of misfolded proteins can not only compromise the integrity of the cellular proteome, but also cell viability. As proteins accumulate and exceed the degradation capacity of the cell, they may be compartmentalized depending on their biochemical properties. There are specific types of compartments (conserved from yeast to mammals) that, according to the solubility state of the proteins, have been designated as juxtanuclear quality control (JUNQ), or insoluble protein deposit (IPOD) [1, 2]. Interestingly, it was demonstrated that the JUNQ compartment participates in an asymmetric segregation of proteins during cell division when these are confined/encaged by the cytoskeletal protein vimentin [3]. This mechanism represents an important strategy for cellular rejuvenation, as observed in single-cell organisms like *Saccharomyces cerevisiae* [3], and is also likely present in stem cells.

Alpha-synuclein (aSyn) is a small protein of 140 amino acids encoded by the *SNCA* gene [4]. aSyn is known as an intrinsically disordered protein mainly found in the cytoplasm and in pre-synaptic terminals, but is also present in the nucleus [5–7]. The function of aSyn is still not entirely clear, but it seems to be related with synaptic vesicle release and biology [8–10].

aSyn is a major component of Lewy bodies (LBs) and Lewy neurites, typical pathological hallmarks of PD and DLB [11]. LBs are typically described as spherical structures composed of filaments of around 10-25 nm in diameter. These filaments radiate out from a dense core (with amorphous material and membranous composition) to the periphery [12–14]. Depending on their localization and morphology, they can be useful for the differential diagnosis of LBD. For example, LBs in cerebral cortex of a patient with DLB typically lack a distinctive core and halo, but they display similar morphologies to LBs from brainstem areas [15, 16].

For a long time, LBs were thought to be the culprits of these disorders. However, several studies demonstrated that LBs are present in surviving neurons of Lewy body disease patients, and can also accumulate in the brains of individuals who display no clinical sign of disease. These and other observations suggest that, possibly, smaller oligomeric forms of aSyn might actually constitute the proteotoxic agents. Therefore, there is a great demand to understand the ultrastructure and architecture of these cytoplasmic inclusions.

One major limitation in the field has been the poor characterization of different types of aSyn assemblies in real biological contexts, such as the cellular environment. With the development of novel microscopy techniques and labeling tools, it is now possible to begin to uncover the molecular architecture of aSyn inclusions and to design experiments to address the correlation between toxicity and different types of assemblies.

Advances in electron microscopy technology have enabled a clear and more accurate characterization of aSyn-immunopositive pathological inclusions. Recent studies showed that LBs have a lamellar structure, where phosphorylated aSyn is at the periphery, and the core is enriched in lipids, rather than proteins [17, 18]. Nevertheless, as exciting as this type of microscopy is, this technology is of rather low throughput, and is not accessible to every laboratory, as it requires special instrumentation and highly-trained specialists. Epifluorescence microscopy, on the other hand, is a standard technique that can be applied in any cell and molecular biology laboratory. However, as any conventional light microscopy approach, it is limited to resolve fine structures due to the diffraction limit of light (∼200-250 nm). The development of expansion microscopy (ExM), a new type of super-resolution microscopy, makes it possible to optically resolve fluorophores that were, otherwise, too close to be distinguished as individual points. This is achieved by anchoring the specimen to a swellable polymer network that can then be used to physically separate the fluorophores, without changing the molecular architecture of the underlying samples [19–21]. The advantage of ExM is that the physical magnification enables higher resolution using a conventional microscope, which makes super-resolution straightforward to implement. Furthermore, it can even be superior to other types of super-resolution imaging techniques such as Stimulated Emission Depletion (STED) microscopy, as shown recently (X10 ExM), with a resolution of around 25 nm [21].

Here, we applied X10 ExM to investigate the molecular architecture of aSyn in cell models, in mice, and in human brain tissue. Using two different models of aSyn inclusion formation, we could distinguish different types of aSyn accumulations. Furthermore, we investigated the spatial organization between aSyn inclusions and intermediate filaments, cytoskeletal components that are present both in cells and in human tissue, and the distribution of aSyn in relation to the vesicular proteins synaptophysin and synapsin I. We observed that aSyn inclusions formed in cell models were surrounded/encaged by vimentin, consistent with our observations in the *cingulate gyrus* from a patient with DLB. This suggests that a structural neurofilament-framework around aSyn deposits might be important for compartmentalization. Interestingly, we found that clusters of aSyn colocalized with synapsin I, a protein known to exist in different types of liquid phases [22].

In summary, X10 ExM enabled us to explore aSyn assemblies and distribution using light microscopy with unprecedented resolution, gaining insight into the architecture and biology of aSyn inclusions. Future studies will enable us to probe and test strategies that interfere with the process of aSyn aggregation.

## Results

### Implementation and workflow of X10 expansion microscopy

To characterize different types of aSyn assemblies formed in relevant biological systems, we assembled a pipeline for applying X10 ExM to cell culture and to mouse and human brain tissue samples (Fig. 1A). To determine the nanoscale resolution achieved, we validated the X10 ExM protocol [21] by estimating the size of the nucleus in HEK cells, prior to and after X10 ExM (Fig. 1C and D). According to the calculations, the expansion factor was around 9,4 times, i.e., consistent with a ~10-fold increase in resolution (more information in the Material and Methods section).

**Figure 1.**
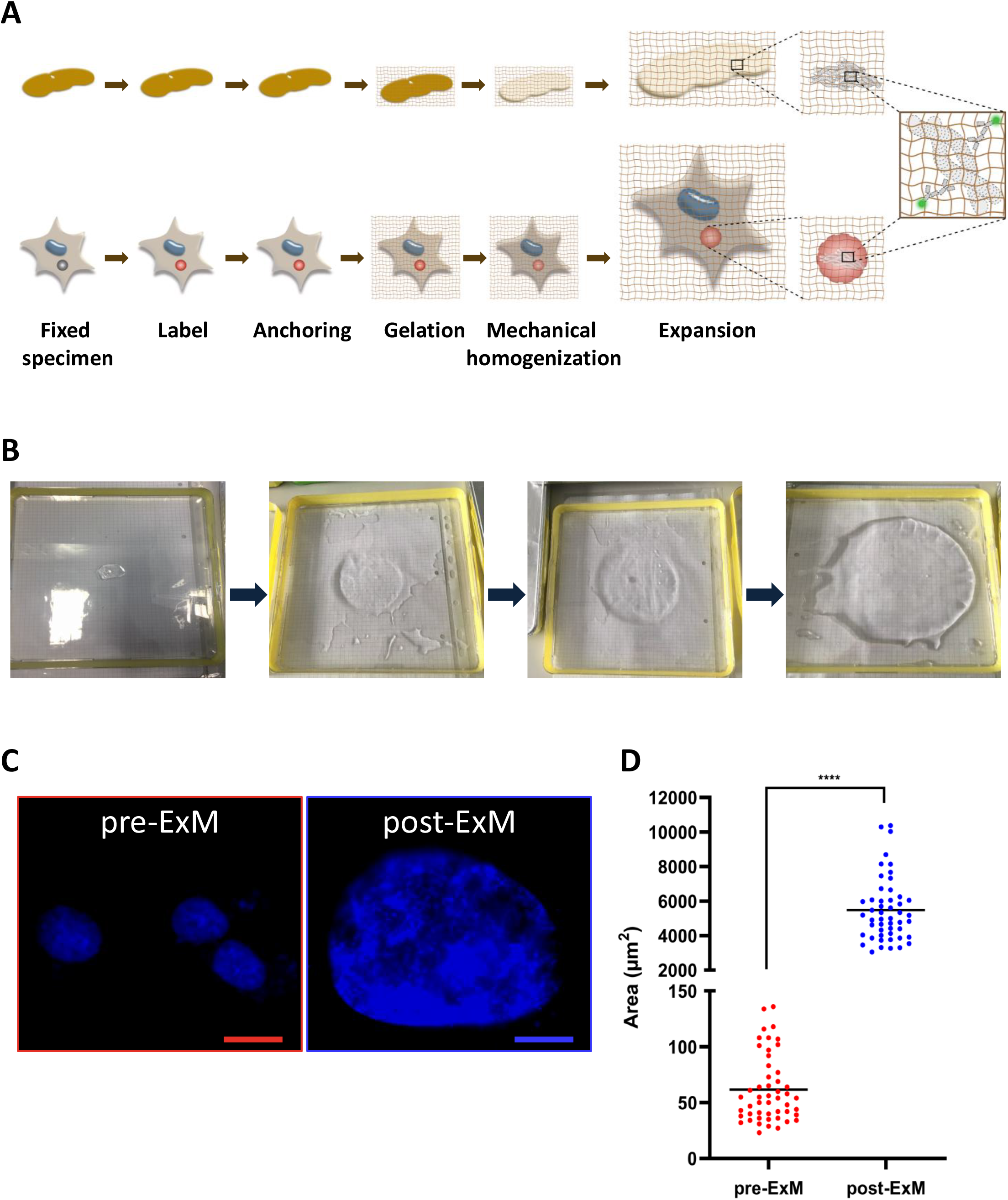
X10 Expansion microscopy workflow and validation. A. Overview of the study. In this study, we investigate the architecture of aSyn in cell models, mice and human tissue using ExM. B. Gel expansion process. Gel expansion was performed by consecutive steps of ddH2O exchanges. C. Exemplary nucleus before and after expansion. Representative images of HEK cells nuclei stained with DAPI were measured to determine the expansion factor. Scale bar: 20μm D. Quantification of expansion factor. The diameter of the nucleus of 50 different cells pre- and post-expansion were evaluated to determine the expansion factor. The average area of pre-ExM is 62μm^2^ (in red) and 5487 μm^2^ post-ExM (in blue). Student’s *t*-test ***p<0.001.

### X10 ExM physically magnifies aSyn assemblies in cells

Next, we investigated the architecture of aSyn assemblies in cells by using two widely used cell models: a model of oligomerization based on the Bimolecular Fluorescence Complementation (BiFC) of Venus fluorescent protein fragments (herein referred to as aSyn BiFC model) [23, 24]; and a model of inclusion formation, based on the co-expression of a C-terminally modified version of aSyn (SynT) and the aSyn-interacting protein synphilin-1 (for simplicity herein referred to as aggregation model) [25]. In this aggregation model, we can model the formation of different types of aSyn inclusions as we previously demonstrated [23]. For comparison, we also employed cells expressing untagged human WT aSyn, which typically does not form visible inclusions in cultured cells. 48h after transfection, we stained the cells with an antibody against aSyn, and performed the X10 ExM protocol.

Comparing the subcellular distribution of these three aSyn variants, we observed a rather homogeneous distribution for untagged WT aSyn post-X10 ExM (Fig. 2A). However, in the aSyn BiFC and in the aggregation models, X10 ExM enabled us to resolve various types of assemblies we could now distinguish with standard epifluorescence or confocal microscopy. In the aSyn BiFC dimerization/oligomerization paradigm, we observed discrete accumulations within the cytoplasm (reflected by the hotspots in the heat maps) compared to what we observed for untagged WT aSyn (Fig. 2B and supplementary Fig. 1A). Since the X10 ExM protocol destroys the aSyn BiFC signal, due to the mechanical digestion (Fig.1A), we used immunostainings, enabling us to compare the signals in the different models. Quantification of the whole image intensity revealed that the pixel intensity was greater for aSyn BiFC than for the WT aSyn (supplementary Fig. 1B).

**Figure 2.**
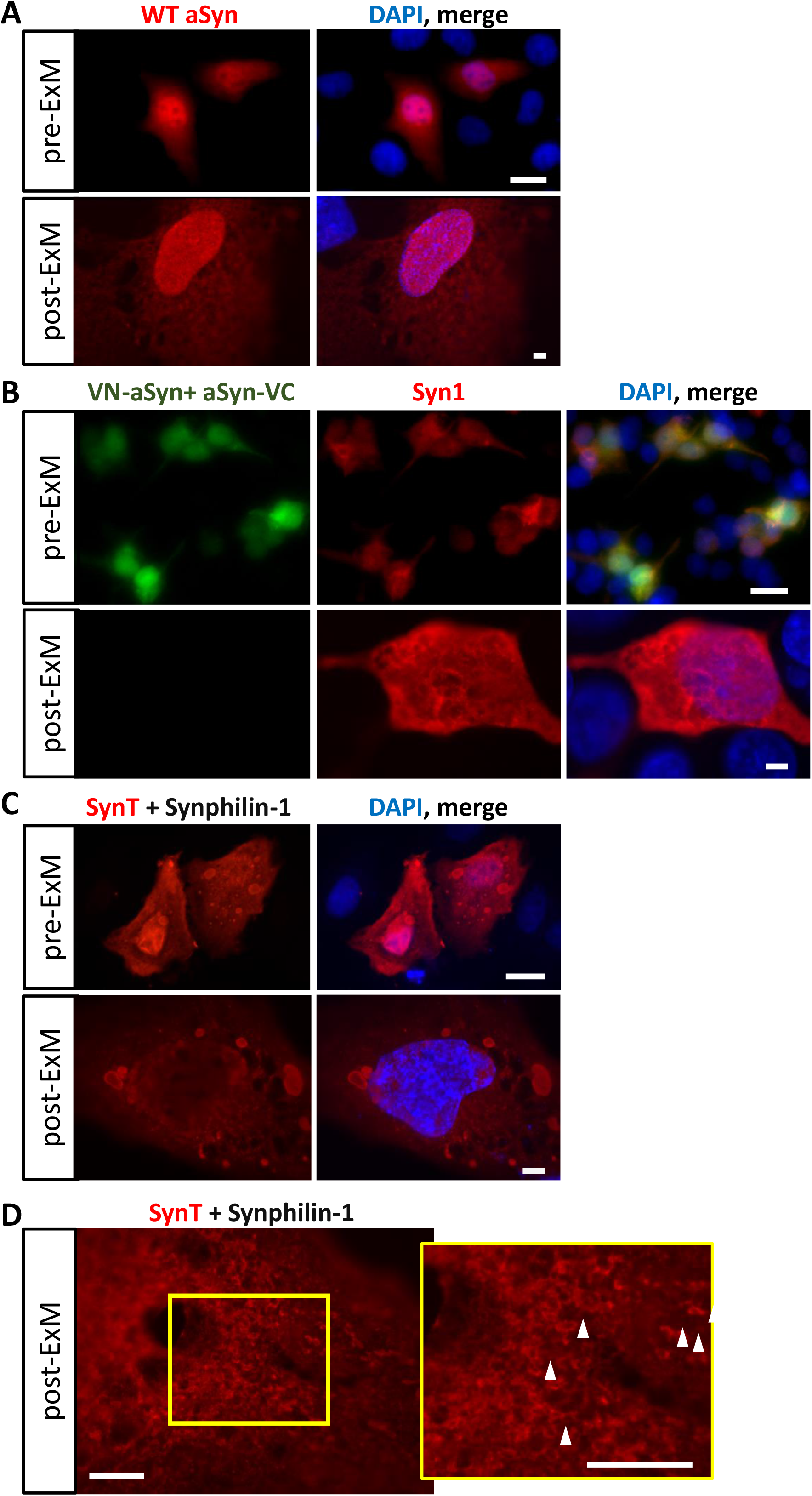
aSyn assemblies in different cell models. Representative images of cells expressing WT aSyn (A), aSyn BiFC model (B), and the aggregation model (C). We observed that aSyn BiFC and aggregation models show various types of aSyn assemblies when compared to those formed by WT aSyn. Scale bar: 20 μm.

Using the aggregation model, we detected different types of aSyn assemblies: inclusions that are typically visible with this model (Fig. 2C), and ring-shaped structures (Fig. 2D) (some of these structures are highlighted with white head arrows). To our knowledge, this is the first report of these aSyn structures and we do not know at this point whether they represent precursor species of the larger inclusions, or other types of accumulations in specific compartments. Overall, the decrowding of biomolecules with ExM allowed us to observe aSyn assemblies at higher resolutions and therefore, to obtain more detailed information about different types of aSyn assemblies in cells.

### Distribution of vimentin in cell lines

Vimentin is an intermediate filament protein type III of particular interest in the context of protein aggregation since it has been implicated in aggresome formation (perinuclear accumulations), and in the elimination of toxic proteins [3]. Recently, it was shown that as aSyn aggregates and acquires a more solid-state, aSyn transforms into perinuclear aggresomes [26].

Resolving intermediate filaments is not trivial, since they fall below the resolution limit of conventional light microscopy. Therefore, we employed X10 ExM and measured several cross-sectional profiles of vimentin in different areas of the cells to determine the number of individual vimentin strands that can be resolved in the two-dimensional (2D) projection (Fig.3A). As expected, we identified more vimentin filaments *per* μm after expansion (Fig.3B). It is important to refer that this analysis refers to the images acquired in this study, since the use of a higher magnification objective might show closer values between pre- and post-expansion. In addition, after X10 ExM, we observed a pearls-on-a string pattern (supplementary data Fig.1C), as previously reported [21], due to incomplete epitope coverage.

**Figure 3.**
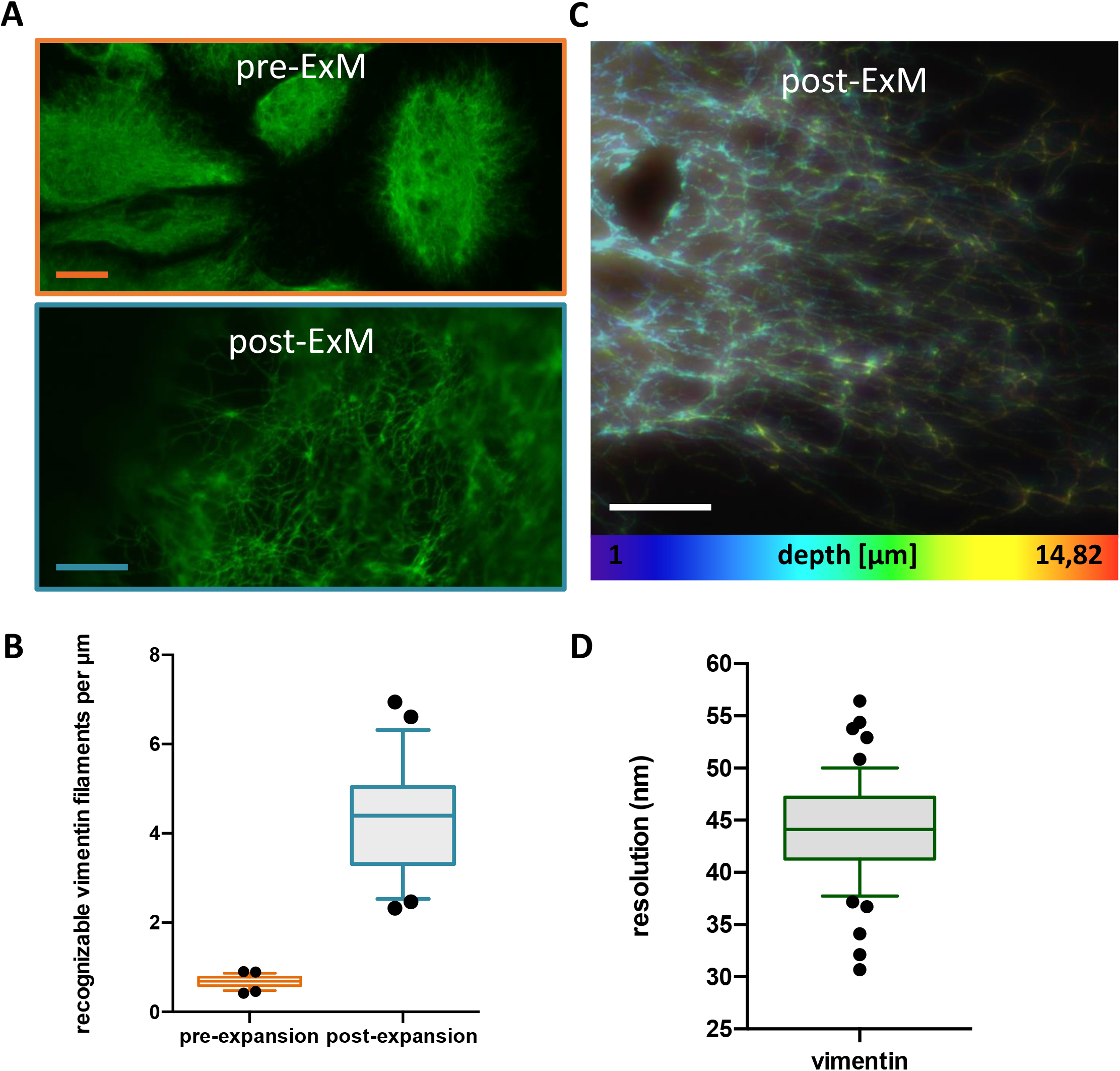
X10 ExM reveals detailed vimentin structure in human cells. A. Typical vimentin morphology observed pre-X10 ExM and post-X10 ExM. B. Different cross-sectional profiles of vimentin were used to determine the number of individual vimentin strands that can be resolved after expansion. 25 line (mean ± SD) scan were analyzed. C. Color-code z-stack of vimentin structure. Projection of 12 z-planes was color coded to represent the ‘depth’ of the stack. D. Analysis of vimentin full-width at half-maximum (FWHM). Vimentin widths yielded average Gaussian-fitted full-width at half-maximum (FWHM) of ± 44 nm (mean ± SD of 59 vimentin profiles). All the scale bars are 20 μm.

Next, individual post-X10 ExM vimentin filaments were measured and fitted with Gaussian functions. From the Full-Width-at-Halve-Maximum (FWHM), we calculated the resolution obtained with X10 ExM (Figure 3D). The overall resolution was around 44 nm, highlighting the power of the technique and its applicability to biological questions.

### Vimentin surrounds aSyn inclusions

The sequestration of misfolded cytosolic proteins into different quality control compartments depends on their aggregation state [1, 2]. Vimentin participates in cell division by establishing mitotic polarity in immortalized mammalian cells [3]. To investigate the relationship between aSyn and vimentin, we started by determining the localization of aSyn inclusions using established markers of specific compartments. Using a misfolded version of the von Hippel-Lindau protein (VHL), a typical marker of the JUNQ quality control compartment, we observed the colocalization of aSyn inclusions with VHL, confirming they can accumulate in this compartment (Fig.4A). Next, we asked, if these inclusions were surrounded by vimentin. We observed that aSyn was surrounded by vimentin (Fig.4B). Using X10 ExM, we dramatically improved the resolution and obtained molecular resolution of the 3D architecture of the inclusions, confirming the presence of vimentin adjacent to aSyn (Fig.4C). These findings shed new light into the molecular mechanisms involved in aSyn-inclusion formation and handling in a cellular context.

**Figure 4.**
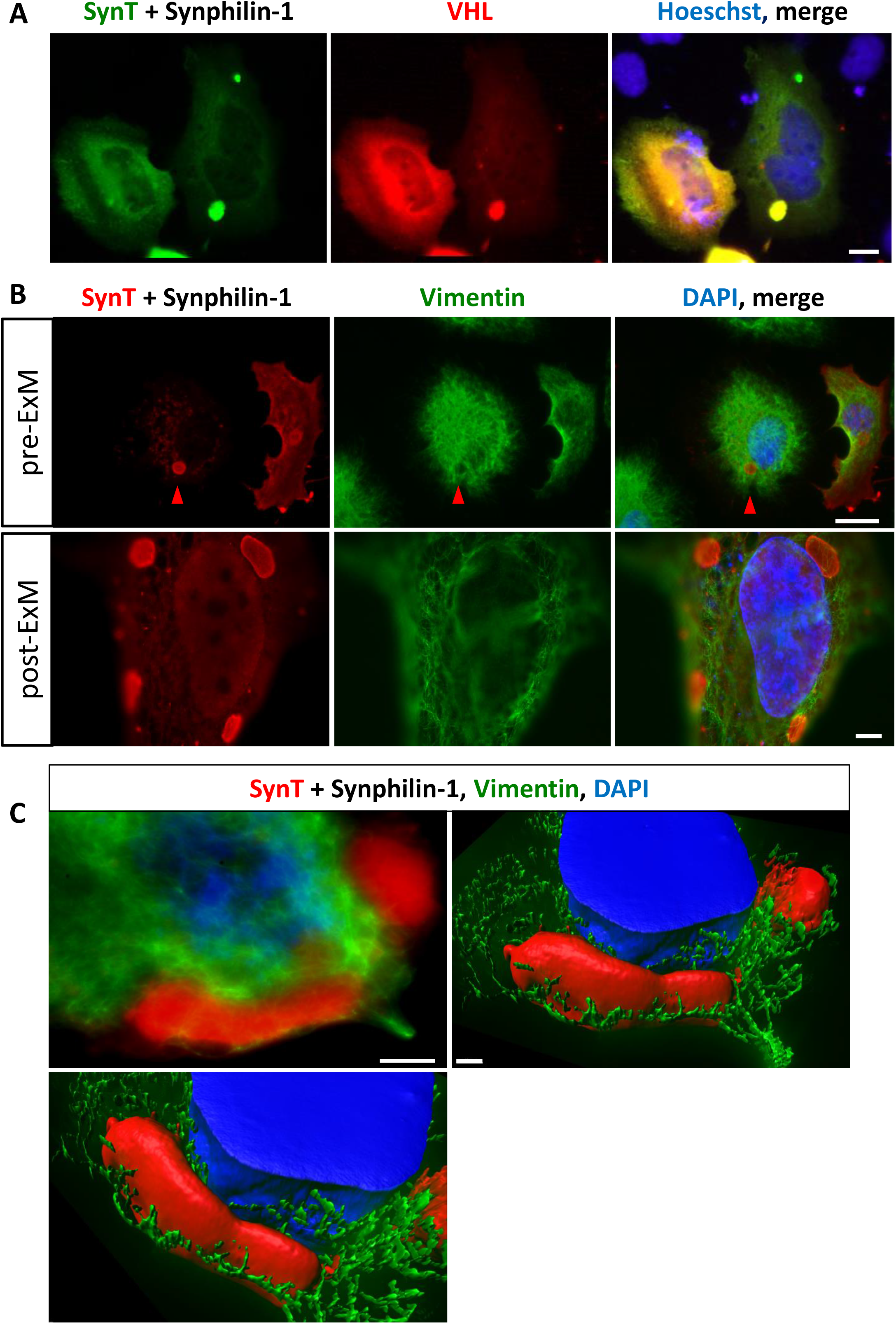
Vimentin surrounds aSyn inclusions. A. aSyn inclusions are present in the JUNQ compartment. Human H4 cells expressing SynT+ Synphilin-1 together with the von Hippel-Lindau protein (VHL) protein, a marker for the JUNQ compartment, shows colocalization of aSyn with VHL. This suggests that aSyn can accumulate at the JUNQ compartment. Scale bars: 10 μm. B-C. aSyn is surrounded/encaged by vimentin. B. Microscopy images from pre- and post-ExM of cell with aSyn inclusions (red) and vimentin (green) shows that aSyn is surrounded/encaged by vimentin. Scale bars: 20μm. C. The top left image shows the sum projection of a representative cell, and the upper right and bottom left image an Imaris 3D reconstruction of the same cell. Scale bar: 10 μm.

### X10 ExM imaging of aSyn in mouse brain tissue

Imaging stained tissue can be a complicated process, as high levels of background are often observed. The complexity and stiffness of the tissue can lead to distortions and aberrations upon expansion. Therefore, it is essential to ensure these are excluded. As proof of concept, we first adapted the protocol to mouse brain tissue, demonstrating its effectiveness for imaging aSyn (Fig. 5). As described previously, we pre-incubated the brain-slices in the monomer solution [27], and adjusted the amount of TEMED [21]. Furthermore, the incubation period was extended up to 48h to ensure successful tissue expansion. In particular, we confirmed the association of aSyn with synaptic vesicle markers, such as synaptophysin, confirming the typical distribution of aSyn in the adult brain (Fig 5B) [28, 29].

**Figure 5.**
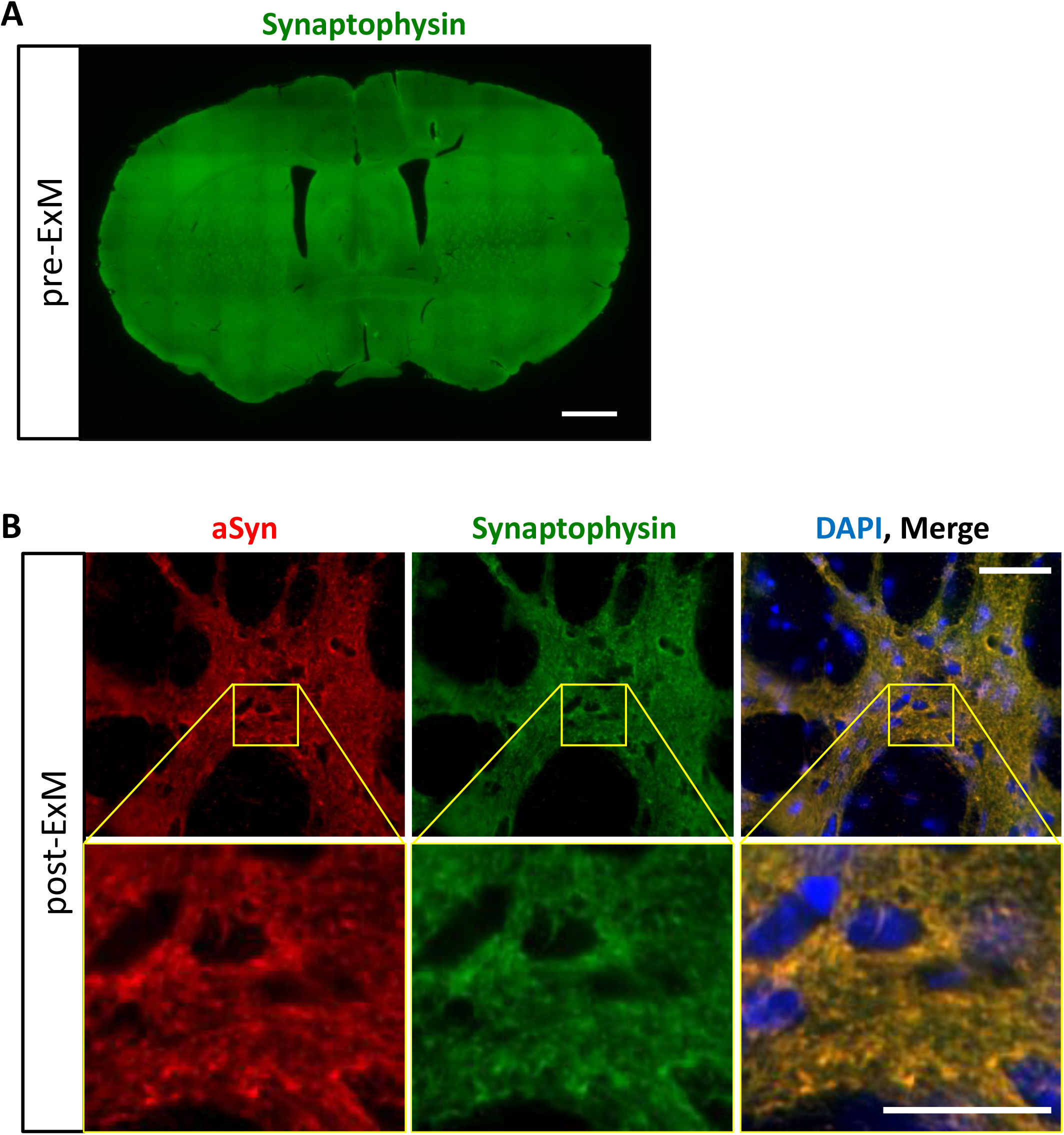
aSyn distribution in the brain of human-WT aSyn transgenic mice. A. Organization of synaptophysin at WT human-aSyn transgenic mice brain. Coronal sections of WT human-aSyn transgenic mice were immunostained for synaptophysin (green). Due to the crowed environment no defined structure is identified. B. Colocalization of aSyn with synaptophysin. Imaging of mouse brain tissue post-ExM shows colocalization of aSyn and synaptophysin. Scale bar: 20μm.

### aSyn co-localizes with synapsin I and is surrounded by neurofilament in DLB brain tissue

Post-mortem neuropathological analyses is still the ultimate approach for confirming and distinguishing between neurodegenerative diseases. However, recent studies show the occurrence of mixed pathology in the brains of patients with neurodegenerative disease and, surprisingly, the occasional occurrence of aSyn Lewy pathology in individuals lacking clinical features of synucleinopathy. Thus, it is essential to develop and apply novel technical approaches for achieving a better characterization of the protein aggregates present in the human brain in order to understand the underlying disease mechanisms. Here, as a proof of concept to characterize pathological aSyn assemblies in the human brain, we analyzed *cingulate gyrus* tissue from a DLB patient. We observed that after X10 ExM, aSyn is observed in multiple small clusters (Fig. 6). In some inlcusions, we also observed aSyn as tubular structures that have not been previously described using conventional microscopy approaches (movie file). Importantly, we also observed that aSyn colocalized with synapsin I (Fig. 6A fluorescence intensity profile), but not with ubiquitin (Fig.6B). This suggests that, in human tissue, aSyn is associated with synaptic vesicles, and that not all aSyn is ubiquitinated, as expected.

**Figure 6.**
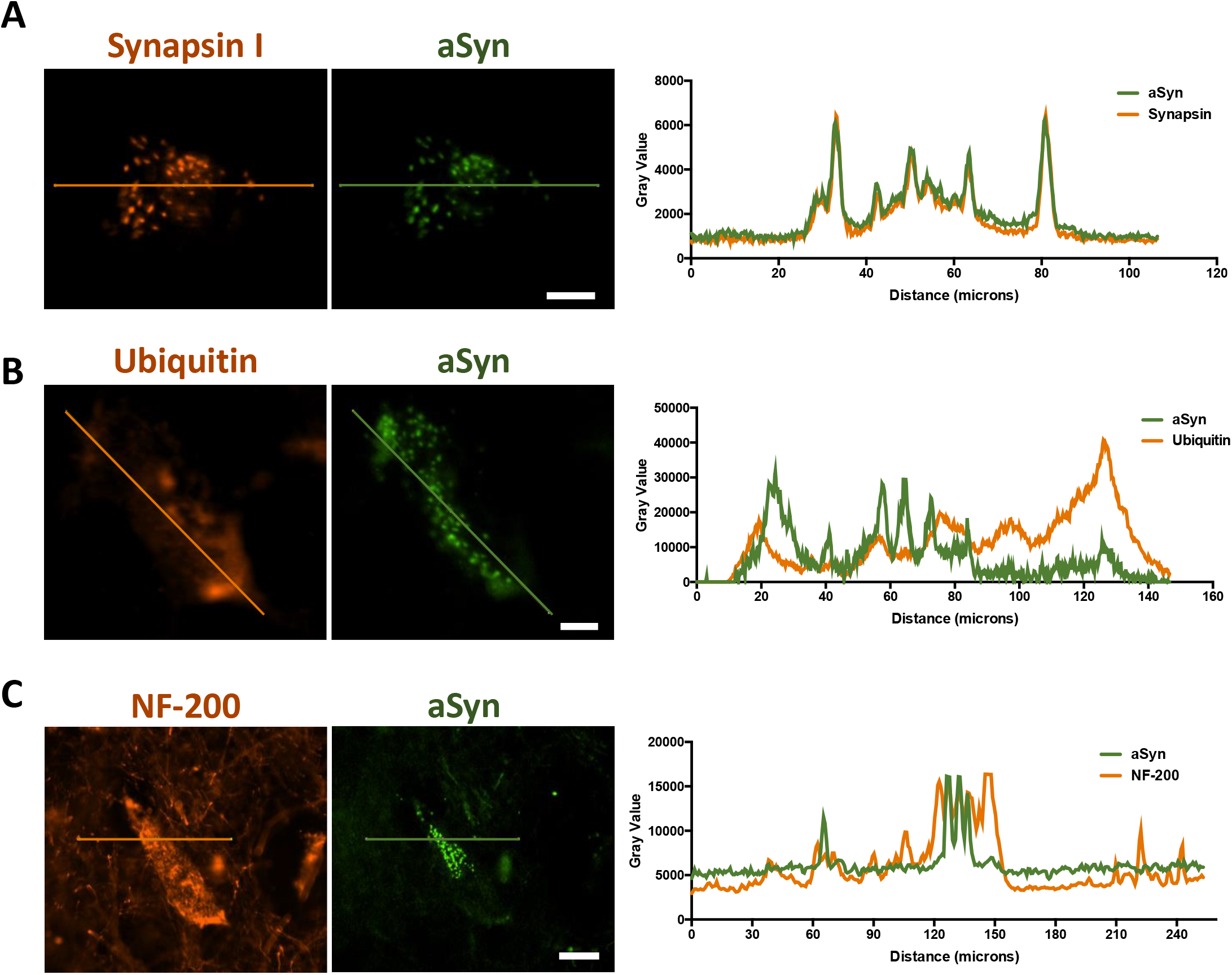
Analysis of aSyn assemblies in DLB tissue. A-B. aSyn colocalizes with synaptic vesicles, but not with ubiquitin. After expansion, aSyn clusters were visualized to show colocalization with synapsin I. This suggests that aSyn associates and/or is within synaptic vesicles. C. Neurofilament surrounds aSyn inclusions. NF-200 surrounds aSyn inclusions, and this is also shown by the line scan, where the aSyn intensity peaks (green) are flanked by NF-200 (orange). Scale bar: 20μm.

Next, we tested the hypothesis that intermediate filament proteins might be involved in the accumulation of aSyn in human brain tissue. In agreement with our observation in cultured cells, with vimentin, we found that aSyn assemblies are surrounded by another intermediated filament protein, neurofilament-200 (NF-200) (Fig. 6C). This was confirmed by the fluorescence intensity profiles, where many aSyn signal peaks (green) were flanked by NF-200 signal (orange) (Fig. 6D, graph). This indicates that these two proteins do not share the same physical space, suggesting that that NF-200 surrounds aSyn.

## Discussion

LBs are the main pathological hallmark of PD and DLB. However, the current hypothesis is that the “smaller” aSyn species, referred to as oligomeric species, and not LBs *per se*, are considered to be the toxic forms. Unfortunately, the term ‘oligomer’ is vague and used loosely in the field, failing to capture the level of detail that would be necessary to define an entity that could be used as a therapeutic target. In addition, we lack adequate methods and tools to characterize the various types of protein assemblies in the complex biological context of a cell and tissue.

ExM is an evolving technique that provides nanoscale precision while preserving biological information. ExM does not require intense computational processing like other super resolution techniques, making it overall an easier technique to be applied even in the context of diagnostics. In fact, this technique has already provided unprecedented knowledge of pathological changes in human tissue [30], or in mapping gene expression sites in the brain [31]. In our study we used X10 ExM to investigate the molecular architecture of aSyn in cells and brain tissue.

We observed that aSyn can accumulate in assemblies with different morphologies, from nanoscale clusters, to ring or even tubular-shaped structures. Furthermore, these depositions can be surrounded by intermediate filament proteins such as vimentin or neurofilament (depending on the biological system). To our knowledge, this is the first report showing this conserved property, from a single cell system to a complex organ as the human brain. At this point, we do not fully understand the biological implications of these assemblies on pathology, and future, dedicated studies with increased numbers of brain samples will be necessary to better address these questions. We speculate that these small assemblies, which are not fully resolved by conventional microscopy, may be early events/steps in the aggregation process, and that tubular-shaped structures surrounded by vimentin may represent aSyn that accumulates over time. We can not rule out that the ring-shaped structures observed may be aSyn in vesicles, like endosomes. Again, additional studies will be necessary to determine the pathological implications of our findings. However, our study establishes the tools and provides the foundation for additional investigations. A novel aspect in our study is the putative role of neurofilaments in aSyn aggregation. Our study suggests that neurofilaments might be important for the formation/accumulation of aSyn. Importantly, we documented similar phenomena both in simple cell models and in human brain tissue. It is possible that intermediate filaments serve as a scaffold for the translocation/sequestration of aSyn into specific areas/compartments of the cells. Considering that aSyn can accumulate in the JUNQ compartment, this suggests a possible biological mechanism for the compartmentalization of specific aSyn assemblies for degradation.

In conclusion, X10 ExM proved to be a powerful imaging technique for studying aSyn accumulation both in cells and tissue. This technique affords not only nanoscale resolution using conventional microscopes, but also facilitates colocalization studies because of the decrowding of the complex biological context.

## Acknowledgements

DFL is supported by the Thiemann-fellowship. TFO is supported by the Deutsche Forschungsgemeinschaft (DFG, German Research Foundation) under Germany’s Excellence Strategy -EXC 2067/1-390729940, and SFB1286 (Project B8).

## Author contributions

PW, DFL, LP, and PIS performed experiments. PW, DFL and TFO designed the study, interpreted data and wrote the manuscript. CS helped with work on human tissue. All authors revised the manuscript.

## Materials and Methods

### Cell culture

Human neuroglioma cells (H4) were maintained in Opti-MEM I Reduced Serum Medium (Life Technologies- Gibco, Carlsbad, CA, USA) and Human Embryonic Kidney 293 (HEK) cells were grown in Dulbecco’s Modified Eagle Medium (DMEM, Life Technologies- Invitrogen, Carlsbad, CA, USA). Both media were supplemented with 10% Fetal Bovine Serum Gold (FBS) (PAA, Cölbe, Germany) and 1% Penicillin-Streptomycin (PAN, Aidenbach, Germany), and kept at 37°C in an atmosphere of 5% CO2. Cells were plated on glass coverslips (18 mm; VWR, cat. no. 631-0153) in a 12-Well cell culture plate (Costar, Corning, New York, USA).

### Cell transfection

HEK cells were transfected with equimolar amounts of VN-aSyn and aSyn-VC encoding plasmids using Metafectene (Biotex, Munich, Germany) as we previously described [23, 24].

H4 cells were transfected with equal amounts of SynT or WT aSyn together with synphilin-1 using FuGENE6 Transfection Reagent (Promega, Madison, USA) in a ratio of 1∶3 according to the manufacturer’s instructions (Lázaro et al. 2014).

### Immunocytochemistry cells

After 48h of transfection, immunocytochemistry was performed. Briefly, cells were fixed with 4% paraformaldehyde for 10 min or 3h (for vimentin stainings) at room temperature (RT). All solutions were prepared in 1x phosphate buffered saline (1xPBS), and incubations were performed at RT if not stated otherwise. After the washes with 1xPBS, the cells were permeabilization with 0.1% Triton X-100 for 20 min. Blocking was performed with 1,5% normal goat serum (NGS) for 1 hour, followed by incubating with primary antibodies overnight (ON) at 4°C (mouse anti-aSyn antibody (211) 1:100 (Santa Cruz Biotechnologies, Santa Cruz, CA, USA); rabbit anti-vimentin 1:100 (Sigma-Aldrich, St. Louis, MO, USA)). Afterwards, cells were washed with 1xPBS three times, and incubated with secondary antibodies at a concentration of 1:100 (Alexa Fluor 488 donkey anti-mouse IgG, Alexa Fluor 568 goat anti-rabbit IgG (Life Technologies-Invitrogen, Carlsbad, CA, USA)) for 2 h. Again, the samples were washed as previously described, and the cells were stained with DAPI at a concentration of 1:1000 for 5 min, and pre-ExM photos were taken.

### Immunohistochemistry staining

All procedures involving animals were performed in accordance with the European Community (Directive 2010/63/EU), institutional and national guidelines (Landesamt für Verbraucherschutz, Braunschweig, Lower Saxony, Germany, license number: 33.9-42502-04-17/2413). All mice were housed at controlled temperature and 12:12 h light/dark cycle.

Mice were sacrificed by exposure to carbon dioxide and afterward perfused transcardially by PFA. After postfixing the brain for 2h at 4°C, they were transferred in 30% of sucrose solution at 4°C till they sank. The brains were stored at −80°C until sectioning.

Brain tissue from a patient with DLB was obtained from the archives of the Institute of Neuropathology, UMG. The cingulate gyrus was dissected, fixed in buffered formalin, and embedded in a mixture of egg albumin, gelatine, sucrose, and glutaraldehyde. 20 μm vibratome sections were cut. Free floating sections were washed in PBS and further processed for ExM. The study on human tissue was approved by the ethics committee of the University Medical Center Göttingen (2/3/19).

For immunohistochemistry on paraffin-embedded human brain tissue, sections of 3 μm thickness were pretreated by heat-induced antigen retrieval in Tris/EDTA buffer pH 8.0 for 30 min in a steamer followed by the incubation with 1:100 anti-aSyn monoclonal antibody (LB509, SIG-39725, Covance, Princeton, NY, USA). Antigen-binding was visualized using an alkaline-phosphatase anti-alkaline phosphatase-based technique using fast red as a chromogen.

For X10 ExM, mouse and human brain slices of 20 μm thickness were subjected to antigen retrieval in 10 mM citrate (Sigma-Aldrich, St. Louis, MO, USA) + 0.5% Tween-20 (Merck, Darmstadt, Germany) at 80°C [32]. Then, the slices were simultaneously blocked and permeabilized in 1xPBS + 2.5% BSA + 0.1% Triton X-100, three times for 10 min, followed by the primary antibodies incubation (for the mouse tissue: mouse anti-aSyn antibody (Syn1) (BD Transduction Laboratory, New Jersey, USA); guinea pig anti-synaptophysin 1 antibody (Synaptic Systems, Göttingen, Germany); and for the human tissue: mouse anti-aSyn monoclonal antibody (LB509, SIG-39725, Covance, Princeton, NY, USA), monoclonal anti-neurofilament-200 (Phosphorylated and Non-Phosphorylated) Clone N52 (Sigma-Aldrich, St. Louis, MO, USA); rabbit anti-synapsin 1 (SYN1) antibody (Synaptic Systems, Göttingen, Germany); rabbit anti-ubiquitin antibody (Abcam ab7780, Cambridge, MA, USA). All antibodies were used at a dilution of 1:100 in the blocking/permeabilization solution, at 4°C ON. The slices were then washed in blocking/permeabilization solution three times for 10 min, and incubated with secondary antibodies prepared in 1:100 dilution in the same blocking/permeabilization solution, for 2h at RT. The following secondary antibodies were used: Alexa Fluor 488 goat anti-rabbit IgG (Life Technologies- Invitrogen, Carlsbad, CA, USA), Alexa Fluor 555 goat anti-rabbit IgG (Life Technologies- Invitrogen, Carlsbad, CA, USA), Alexa Fluor 488 goat anti-mouse IgG (Life Technologies- Invitrogen, Carlsbad, CA, USA), Alexa Fluor 555 donkey anti-mouse IgG (Life Technologies-Invitrogen, Carlsbad, CA, USA), Alexa Fluor 546 goat anti-guinea Pig IgG ((Life Technologies-Invitrogen, Carlsbad, CA, USA). The brain slices were again washed three times for 10 min as before, and three times for 10 min in high-salt PBS (PBS + 350 mM NaCl), followed by two times for 10 min in PBS.

### X10 ExM procedure for cultured cells

The expansion protocol was performed as previously described [32]. Briefly, after immunohistochemistry, the samples were incubated with 500μl of anchoring buffer (1x PBS + 0.1mg/ml Acryloyl-X (Life Technologies, Carlsbad, CA, USA)) ON at RT. Then, the samples were washed with 1x PBS three times for 5 min, and the gelation solution was applied.

For the X10 gelling solution, 33% (w/w) of high purity sodium acrylate (SA) and *N,N*-dimethylacrylamide acid (DMAA) monomers solution was dissolved at a molar ratio of 4:1 (DMAA:SA) in ddH_2_O. Then, the gelling solution was bubbled with nitrogen gas for 40 min at RT. In a separated tube, potassium persulfate (KPS) (0.036 g/ml stock of KPS) was prepared at 0.4 molar% relative to the monomer concentration, and mixed with the DMAA+SA solution. Another 20 min of bubbling with nitrogen gas was performed in a mix of ice + water. For 5ml of gelling solution, 20μl of *N*,*N*,*N*′,*N*′-tetramethyl-ethane-1,2-diamine (TEMED) was added. For cells, 80μl of the solution was applied onto parafilm, and the coverslips were carefully placed on upside down.

### X10 ExM procedure for brain slices

For brain slices, a pre-incubation was performed with the monomer solution + KPS for 10 min on ice, before placing the brain slices in a custom gelation chamber as previously described[27]. Also, we doubled the amount of TEMED added (40 μl for 5 ml gel solution) to the monomer solution. The polymerization of the gel was performed at least 36h up to 48h. Then, the samples were placed into a 12-well plate with 1,5ml of digestion buffer (50 mM Tris buffer, 0.8 M guanidinium chloride, 8 U/ml proteinase K, 0.5% Triton X-100, pH 8.0; all chemicals are from Sigma-Aldrich, St. Louis, MO, USA), and incubated ON at 50°C in a humified chamber.

For the expansion step, the digested gels were placed in a tray with an excess volume of ddH_2_O (at least 10-times the final gel volume), replacing the ddH_2_O every hour for 4-5 times. The final expansion step was done ON.

### Imaging of the X10 ExM gels

For the cell models, image acquisition was performed on a Leica DMI6000B microscope (Leica Microsystems) with a 40x 0.60 numerical aperture (NA) and 63x, 0.70 NA HC PL both air Fluotar objectives (Leica Microsystems).

The imaging of tissue was performed on a Nikon Ti-E epifluorescence microscope (Nikon Corporation, Chiyoda, Tokyo, Japan) with a 10x 0.45 NA air Plan Apochromat objective, a 40x 0.95 NA air Plan Apochromat objective, a Nikon DS-Qi2 camera (Nikon Corporation, Chiyoda, Tokyo, Japan) and operated with the NIS-Elements AR software (Nikon Corporation, Chiyoda, Tokyo, Japan).

Brightfield microphotographs of tissue sections were acquired using a light microscope (BX51, Olympus, Tokyo, Japan) equipped with a digital camera (Dp71, Software CellSens Dimension v.1.7.1, Olympus, Tokyo, Japan).

### Estimation of the expansion factor

To estimate the expansion factor, the size of 50 nuclei of HEK 293 cells was measured before and after expansion, and the calculated factor was achieved according to the formula below.

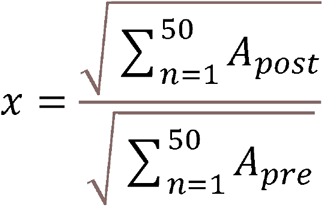

### Imaging and data analyses

The imaging data were analyzed using customized scripts in ImageJ software (Wayne Rasband, NIH, Bethesda, MD, USA), and Imaris (Bitplane) was used to rendering the 3D images. GraphPad Prism 5 software (San Diego California, USA) was used to performed statistic analysis.

### Heat maps

The heat maps were generated to enable a better visualization/representation of aSyn clusters. A custom-written script was created in ImageJ to perform the analysis. Briefly, the aSyn channel was selected, and the z-stacks converted to a maximum intensity projection image. The Fire LUT was applied, and the area of the all image was selected, and measured to obtain the fluorescence intensity. The values were normalized and plotted in the GraphPad. A Student’s *t*-test analysis was performed.

### Fluorescence intensity profiles

The fluorescence intensity profile was measured in ImageJ, by doing a thin cross-section over the target regions (Fig.3A-B and Fig.6). The graphs were generated in GraphPad.

### Z-projection by color-code

To preserve the z-information in a 2D image, the z-stack was color code to represent the ‘depth’ of the stack.

### Determination of vimentin resolution

The resolution was determined by measuring the full width at half maximum (FWHM) of vimentin (Fig. 3D). Briefly, 59 vimentin structures were analyzed, and a custom-written script in ImageJ used to obtain the FWHM. The obtained value was then divided by the expansion factor.

---

**Supplementary Figure 1.**
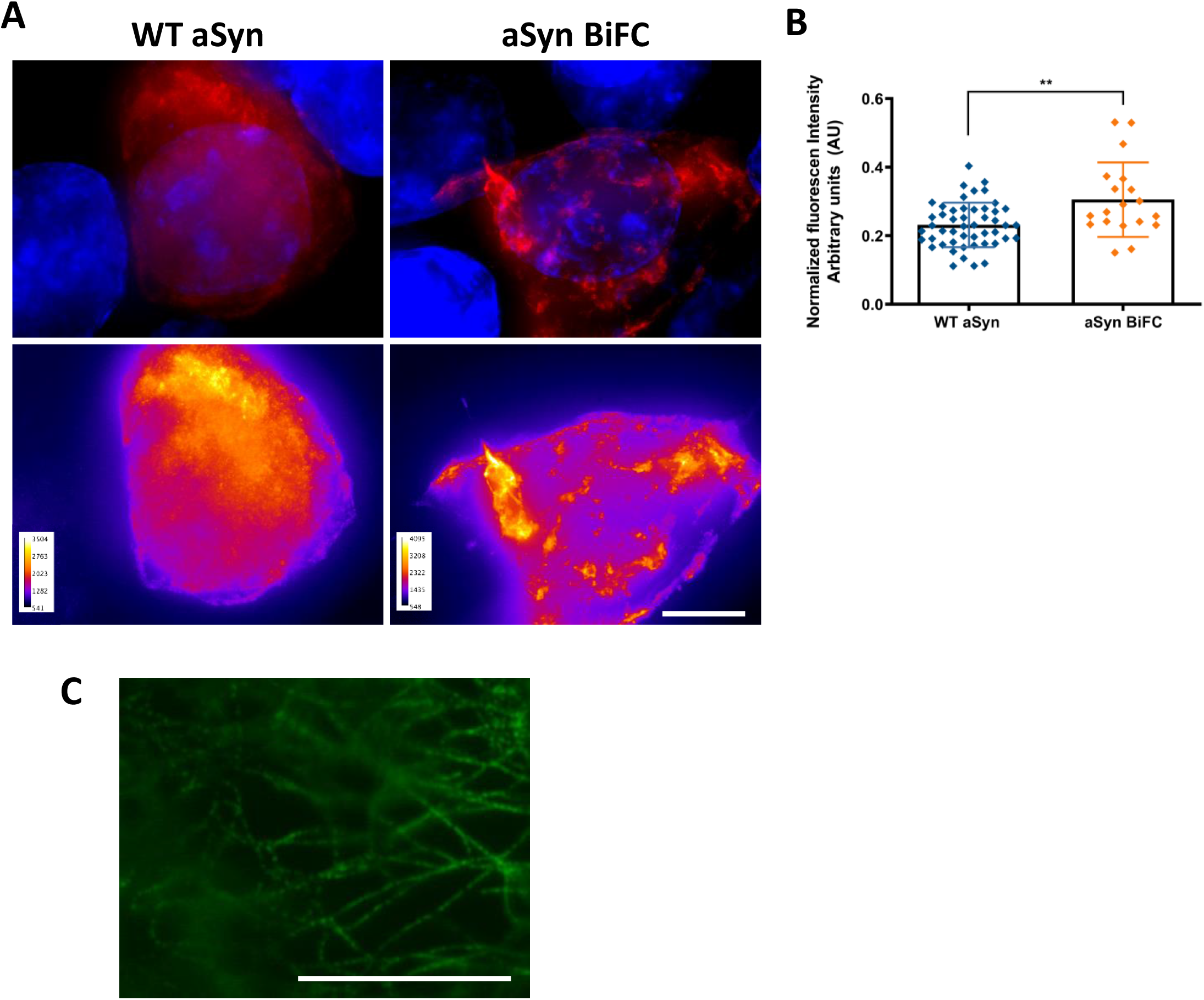
aSyn assemblies in cultured cells. A-B aSyn clusters in the cells. A. Heat maps where generated to better visualize aSyn BiFC clustering in the cells post-ExM. B. The area of the all image was selected, measured and plotted to obtain the fluorescence intensity. Student’s *t*-test **p<0.01. Scale bar: 50μm. C. Vimentin uneven epitope coverage. ExM revels the limitation of using traditional antibodies, reveling vimentin pattern as a pearls-on-a string.

**Supplementary figure 2.**
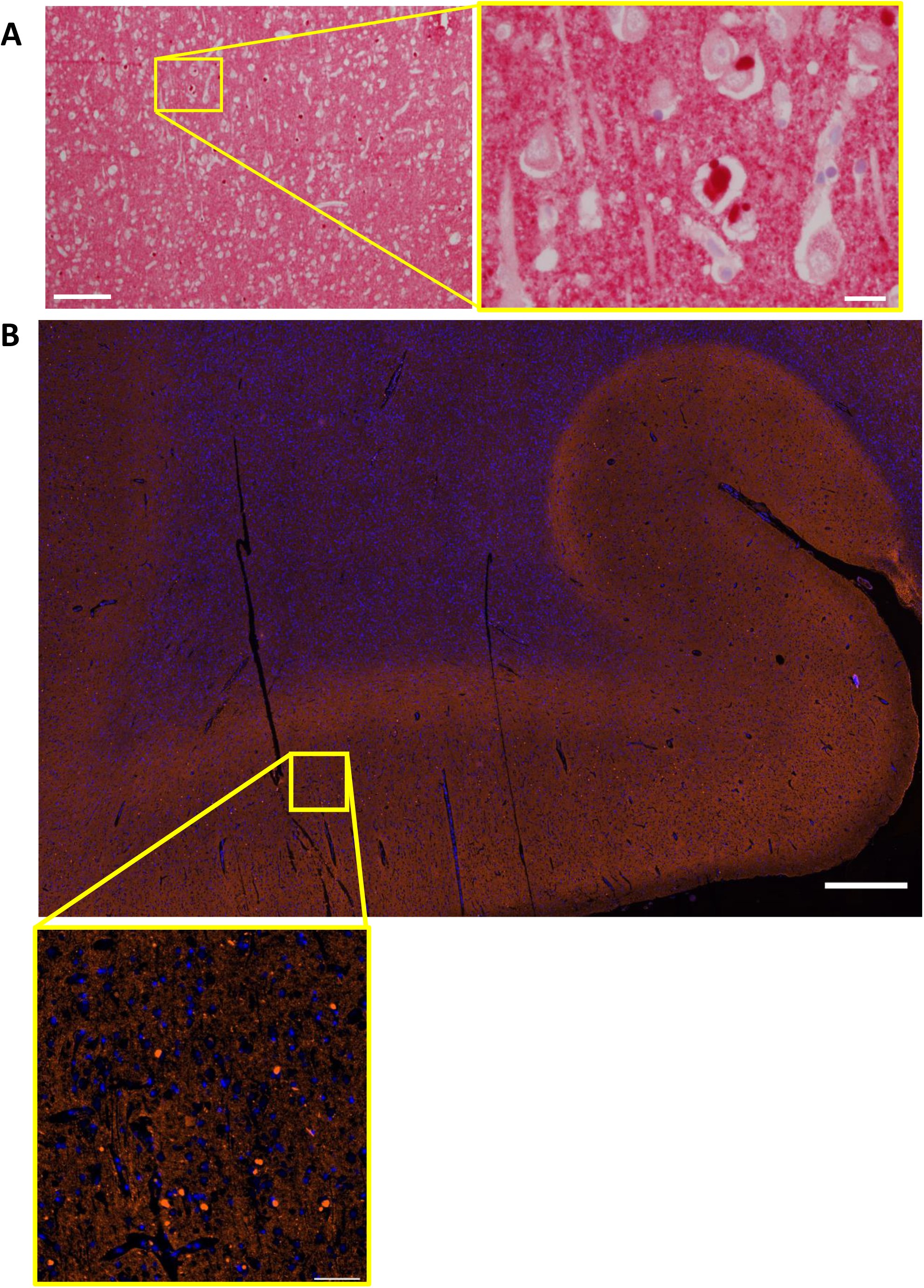
LBs present in DLB brain tissue. **A** LBs immunostained for aSyn (Scale bar left image: 200μm, right image: 20μm). **B.** Coronal section of the human brain tissue section from *cingulate gyrus* stained for aSyn (red) and counter-stained with DAPI (blue) to stain nuclei. Scale bar upper panel: 1000μm; lower panel: 100μm.

**Movie 1. Z-stack of tubular aSyn structures in DLB tissue.** aSyn (green), surrounded by NF-200 (red), can form elongated/tubular structures in human tissue.

## References

1. Kaganovich, D., R. Kopito, and J. Frydman, Misfolded proteins partition between two distinct quality control compartments. Nature, 2008. 454(7208): p. 1088–95.

2. Weisberg, S.J., et al., Compartmentalization of superoxide dismutase 1 (SOD1G93A) aggregates determines their toxicity. Proc Natl Acad Sci U S A, 2012. 109(39): p. 15811–6.

3. Ogrodnik, M., et al., Dynamic JUNQ inclusion bodies are asymmetrically inherited in mammalian cell lines through the asymmetric partitioning of vimentin. Proc Natl Acad Sci U S A, 2014. 111(22): p. 8049–54.

4. Bendor, J.T., T.P. Logan, and R.H. Edwards, The function of α-synuclein. Neuron, 2013. 79(6): p. 1044–66.

5. Maroteaux, L., J.T. Campanelli, and R.H. Scheller, Synuclein: a neuron-specific protein localized to the nucleus and presynaptic nerve terminal. J Neurosci, 1988. 8(8): p. 2804–15.

6. Theillet, F.-X., et al., Structural disorder of monomeric α-synuclein persists in mammalian cells. Nature, 2016. 530(7588): p. 45–50.

7. Pinho, R., et al., Nuclear localization and phosphorylation modulate pathological effects of alpha-synuclein. Hum Mol Genet, 2019. 28(1): p. 31–50.

8. Burre, J., et al., Alpha-synuclein promotes SNARE-complex assembly in vivo and in vitro. Science, 2010. 329(5999): p. 1663–7.

9. Liu, S., et al., alpha-Synuclein produces a long-lasting increase in neurotransmitter release. EMBO J, 2004. 23(22): p. 4506–16.

10. Burre, J., M. Sharma, and T.C. Sudhof, alpha-Synuclein assembles into higher-order multimers upon membrane binding to promote SNARE complex formation. Proc Natl Acad Sci U S A, 2014. 111(40): p. E4274–83.

11. Spillantini, M.G., et al., α-Synuclein in Lewy bodies. Nature, 1997. 388(6645): p. 839–840.

12. Roy, S. and L. Wolman, Ultrastructural observations in Parkinsonism. J Pathol, 1969. 99(1): p. 39–44.

13. Spillantini, M.G., et al., alpha-Synuclein in filamentous inclusions of Lewy bodies from Parkinson’s disease and dementia with lewy bodies. Proc Natl Acad Sci U S A, 1998. 95(11): p. 6469–73.

14. Arima, K., et al., Immunoelectron-microscopic demonstration of NACP/alpha-synuclein-epitopes on the filamentous component of Lewy bodies in Parkinson’s disease and in dementia with Lewy bodies. Brain Res, 1998. 808(1): p. 93–100.

15. Kosaka, K., et al., Presenile dementia with Alzheimer-, Pick- and Lewy-body changes. Acta Neuropathol, 1976. 36(3): p. 221–33.

16. Okazaki, H., L.E. Lipkin, and S.M. Aronson, Diffuse intracytoplasmic ganglionic inclusions (Lewy type) associated with progressive dementia and quadriparesis in flexion. J Neuropathol Exp Neurol, 1961. 20: p. 237–44.

17. Shahmoradian, S.H., et al., Lewy pathology in Parkinson’s disease consists of crowded organelles and lipid membranes. Nat Neurosci, 2019. 22(7): p. 1099–1109.

18. Moors, T.E., et al., Detailed structural orchestration of Lewy pathology in Parkinson’s disease as revealed by 3D multicolor STED microscopy. BioRxiv, 2018.

19. Chen, F., P.W. Tillberg, and E.S. Boyden, Optical imaging. Expansion microscopy. Science, 2015. 347(6221): p. 543–8.

20. Chozinski, T.J., et al., Expansion microscopy with conventional antibodies and fluorescent proteins. Nat Methods, 2016. 13(6): p. 485–8.

21. Truckenbrodt, S., et al., X10 expansion microscopy enables 25-nm resolution on conventional microscopes. EMBO Rep, 2018. 19(9).

22. Milovanovic, D., et al., A liquid phase of synapsin and lipid vesicles. Science, 2018. 361(6402): p. 604–607.

23. Lazaro, D.F., et al., Systematic comparison of the effects of alpha-synuclein mutations on its oligomerization and aggregation. PLoS Genet, 2014. 10(11): p. e1004741.

24. Outeiro, T.F., et al., Formation of toxic oligomeric alpha-synuclein species in living cells. PLoS One, 2008. 3(4): p. e1867.

25. McLean, P.J., H. Kawamata, and B.T. Hyman, Alpha-synuclein-enhanced green fluorescent protein fusion proteins form proteasome sensitive inclusions in primary neurons. Neuroscience, 2001. 104(3): p. 901–12.

26. Ray, S., et al., alpha-Synuclein aggregation nucleates through liquid-liquid phase separation. Nat Chem, 2020. 12(8): p. 705–716.

27. Chen, F., P.W. Tillberg, and E.S. Boyden, Expansion microscopy. Science, 2015. 347(6221): p. 543–548.

28. Reyes, J.F., et al., A cell culture model for monitoring α-synuclein cell-to-cell transfer. Neurobiol Dis, 2015. 77: p. 266–75.

29. Danzer, K.M., et al., Exosomal cell-to-cell transmission of alpha synuclein oligomers. Mol Neurodegener, 2012. 7: p. 42.

30. Zhao, Y., et al., Nanoscale imaging of clinical specimens using pathology-optimized expansion microscopy. Nat Biotechnol, 2017. 35(8): p. 757–764.

31. Martinez, C.C., et al., Spatial transcriptional signatures define margin morphogenesis along the proximal-distal and medio-lateral axes in tomato (Solanum lycopersicum) leaves. Plant Cell, 2021. 33(1): p. 44–65.

32. Truckenbrodt, S., et al., A practical guide to optimization in X10 expansion microscopy. Nat Protoc, 2019. 14(3): p. 832–863.

